# Deep Neural Networks in Computational Neuroscience

**DOI:** 10.1101/133504

**Authors:** Tim C Kietzmann, Patrick McClure, Nikolaus Kriegeskorte

## Abstract

The goal of computational neuroscience is to find mechanistic explanations of how the nervous system processes information to give rise to cognitive function and behaviour. At the heart of the field are its models, i.e. mathematical and computational descriptions of the system being studied, which map sensory stimuli to neural responses and/or neural to behavioural responses. These models range from simple to complex. Recently, deep neural networks (DNNs) have come to dominate several domains of artificial intelligence (AI). As the term “neural network” suggests, these models are inspired by biological brains. However, current DNNs neglect many details of biological neural networks. These simplifications contribute to their computational efficiency, enabling them to perform complex feats of intelligence, ranging from perceptual (e.g. visual object and auditory speech recognition) to cognitive tasks (e.g. machine translation), and on to motor control (e.g. playing computer games or controlling a robot arm). In addition to their ability to model complex intelligent behaviours, DNNs excel at predicting neural responses to novel sensory stimuli with accuracies well beyond any other currently available model type. DNNs can have millions of parameters, which are required to capture the domain knowledge needed for successful task performance. Contrary to the intuition that this renders them into impenetrable black boxes, the computational properties of the network units are the result of four directly manipulable elements: *input statistics, network structure, functional objective*, and *learning algorithm*. With full access to the activity and connectivity of all units, advanced visualization techniques, and analytic tools to map network representations to neural data, DNNs represent a powerful framework for building task-performing models and will drive substantial insights in computational neuroscience.

## Explaining brain information processing requires complex, task performing models

The goal of computational neuroscience is to find mechanistic explanations for how the nervous system processes information to support cognitive function as well as adaptive behaviour. Computational models, i.e. mathematical and computational descriptions of component systems, aim to capture the mapping of sensory input to neural responses and furthermore to explain representational transformations, neuronal dynamics, and the way the brain controls behaviour. The overarching challenge is therefore to define models that explain neural measurements as well as complex adaptive behaviour. Historically, computational neuroscientists have had successes with shallow, linear-nonlinear “tuning” models used to predict lower-level sensory processing. Yet, the brain is a deep recurrent neural network that exploits multistage non-linear transformations and complex dynamics. It therefore seems inevitable that computational neuroscience will come to rely increasingly on complex models, likely from the family of deep recurrent neural networks. The need for multiple stages of nonlinear computation has long been appreciated in the domain of vision, by both experimentalists (Hubel & Wiesel, 1959) and theorists (Fukushima, 1980; Lecun & Bengio, 1995; Riesenhuber & Poggio, 1999; G. Wallis & Rolls, 1997).

The traditional focus on shallow models was motivated both by the desire for simple explanations and by the difficulty of fitting complex models. Hand-crafted features, which laid the basis of modern computational neuroscience (Jones & Palmer, 1987), do not carry us beyond restricted lower-level tuning functions. As an alternative approach, researchers started directly using neural data to fit model parameters (Dumoulin & Wandell, 2008; M. C.-K. Wu, David, & Gallant, 2006). This approach was shown to be particularly successful for early visual processes (Cadena et al., 2017; Gao & Ganguli, 2015). Despite its elegance, importance, and success, this approach is ultimately limited by the amount of neural observations that can be collected from a given system. Even with neural measurement technology advancing rapidly (multi-site array recordings, two-photon imaging, or neuropixels, to name just a few), the amount of recordable data may not provide enough constraints to fit realistically complex, i.e. parameter-rich models. For instance, while researchers can now record separately from hundreds of individual neurons, and the number of stimuli used may approach 10,000, the numbers of parameters in deep neural networks (DNNs) are many orders of magnitude larger. For instance, the influential object recognition network “AlexNet” has 60 million parameters (Krizhevsky, Sutskever, & Hinton, 2012), a more recent object recognition network, VGG-16, has 138 million parameters (Simonyan & Zisserman, 2015). This high number is required to encode substantial domain knowledge, which is required for intelligent behaviour. Transferring this information into the model through the bottleneck of neural measurements alone is likely too inefficient for understanding and performing real-world tasks.

In search for a solution to this conundrum, the key insight was the idea that rather than fitting parameters based on neural observations, models could instead be trained to perform relevant behaviour in the real world. This approach brings machine learning to bear on models for computational neuroscience, enabling researchers to constrain the model parameters via task training. In the domain of vision, for instance, category-labelled sets of training images can easily be assembled using web-based technologies, and the amount of available data can therefore be expanded more easily than for measurements of neural activity. Of course, different models trained to perform a relevant task (such as object recognition, if one tried to understand computations in the primate ventral stream) might differ in their ability to explain neural data. Testing which model architectures, input statistics, and learning objectives yield the best predictions of neural activity in novel experimental conditions (e.g. a set of images that has not been used in fitting the parameters) is a thus a powerful technique to learn about the computational mechanisms that might underlie the neural responses. Together, the combined use of task training- and neural data enables us to build complex models with extensive knowledge about the world in order to explain how biological brains implement cognitive function.

## Brain-inspired neural network models are revolutionising artificial intelligence and exhibit rich potential for computational neuroscience

Neural network models have become a central class of models in machine learning (Figure 1). Driven to optimize task-performance, researchers developed and improved model architectures, hardware and training schemes that eventually led to today’s high-performance DNNs. These models have revolutionised several domains of AI (LeCun, Bengio, & Hinton, 2015). Starting with the seminal work by Krizhevsky et al (2012), who won the ImageNet competition in visual object recognition by a large margin, deep neural networks now dominate computer vision (He, Zhang, Ren, & Sun, 2016; Simonyan & Zisserman, 2015; Szegedy et al., 2015), and drove reinforcement learning (Lange & Riedmiller, 2010; Mnih et al., 2015), speech-recognition (Sak, Senior, & Beaufays, 2014), machine translation (Sutskever, Vinyals, & Le, 2014; Y. Wu et al., 2016), and many other domains to unprecedented performance levels. In terms of visual processing, deep convolutional, feed-forward networks (CNNs) now achieve human-level classification performance (VanRullen, 2017).

**Figure 1.**
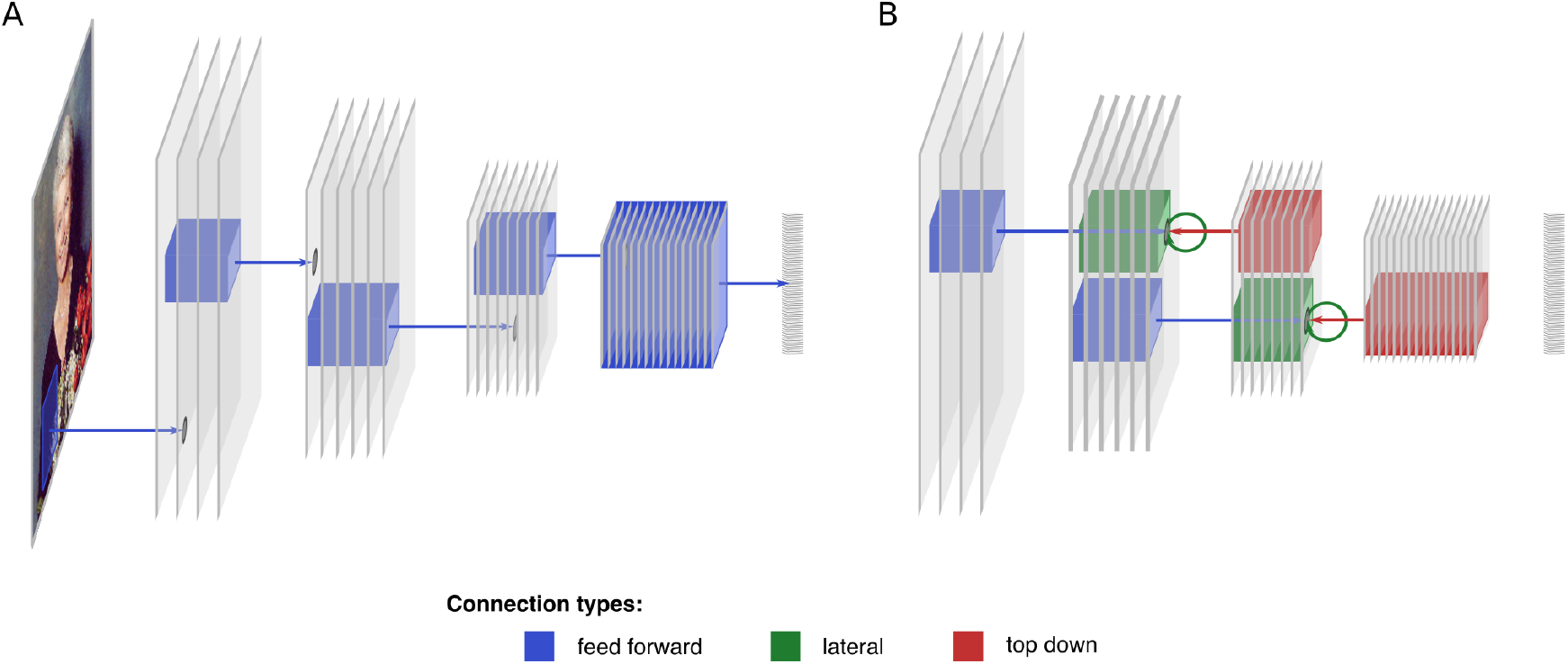
Convolutional neural network structure. (A) An example feed forward convolutional neural network (CNN) with 3 convolutional layers followed by a fully-connected layer. Bottom-up receptive fields for selected neurons are illustrated with blue boxes. (B) The bottom-up (blue), lateral (green), and top-down (red) receptive fields for two example neurons in different layers of a recurrent convolutional neural network (RCNN).

Although originally inspired by biology, current DNNs implement only the most essential features of biological neural networks. They are composed of simple units that typically compute a linear combination of their inputs and pass the result through a static nonlinearity (e.g. setting negative values to zero). Similar to the ventral stream in the brain, convolutional neural networks process images through a sequence of visuotopic representations: each unit “sees” a restricted local region of the map in the previous layer (its receptive field), and similar feature detectors exist across spatial locations (although this is only approximately true in the primate brain). Along the hierarchy, CNNs and brains furthermore perform a deep cascade of non-linear computations, resulting in receptive fields that increase in size, invariance, and complexity. Beyond these similarities, DNNs do typically not include many biological details. For instance, they often do not include lateral or top-down connections, and compute continuous outputs (real numbers that could be interpreted as firing rates) rather than spikes. The list of features of biological neural networks not captured by these models is endless.

Yet, despite large differences and many biological features missing, deep convolutional neural networks predict functional signatures of primate visual processing across multiple hierarchical levels at unprecedented accuracy. Trained to recognise objects, they develop V1-like receptive fields in early layers, and are predictive of single cell recordings in macaque IT (Cadieu et al., 2014; Khaligh-Razavi & Kriegeskorte, 2014; for reviews see Kriegeskorte, 2015; Yamins et al., 2014; Yamins & DiCarlo, 2016; Figure 2A). In particular, the explanatory power of DNNs is on par with the performance of linear prediction based on an independent set of IT neurons and exceeds linear predictions based directly on the category labels on which the networks were trained (Yamins et al., 2014). DNNs explain about 50% of the variance of windowed spike counts in IT across individual images (Yamins et al., 2014), a performance level comparable to that achieved with Gabor models in V1 (Olshausen & Field, 2005). DNNs thereby constitute the only model class in computational neuroscience that is capable of predicting responses to novel images in IT with reasonable accuracy. DNN modelling has also been shown to improve predictions of intermediate representations in area V4 over alternative models (Yamins & DiCarlo, 2016). This indicates that, in order to solve the task of object classification, the trained network passes information through a similar sequence of intermediate representations as the primate brain.

**Figure 2.**
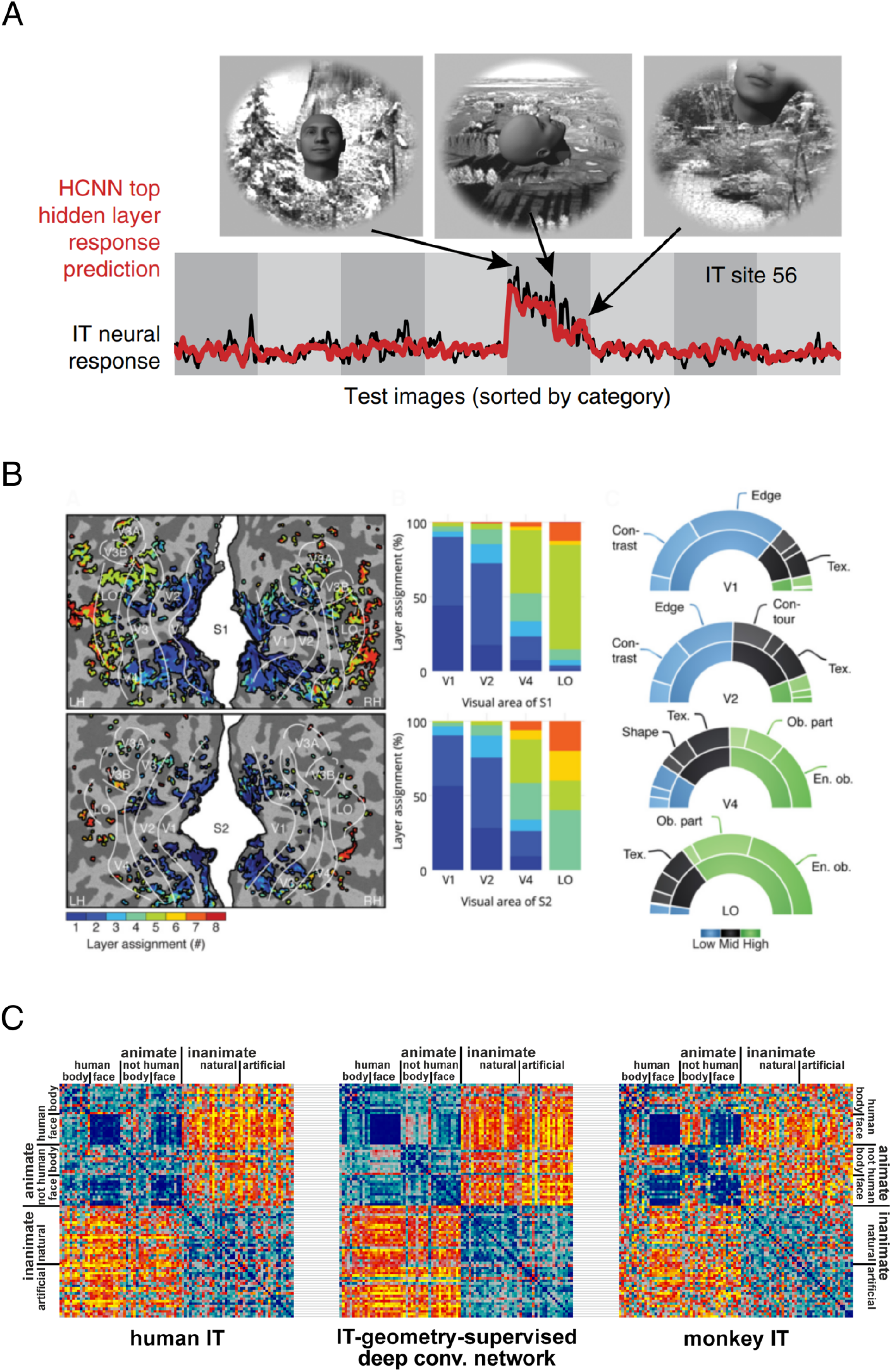
Testing the internal representations of DNNs against neural data. (A) An example of neuron-level encoding with a convolutional neural network (adapted from Yamins & DiCarlo, 2016). (B) A CNN based encoding model applied to human fMRI data (adapted from Güçlü & van Gerven, 2015). (C) Comparing the representational geometry of a trained CNN to human and monkey brain activation patterns using representation-level similarity analysis (adapted from Khaligh-Razavi & Kriegeskorte, 2014).

In human neuroscience too, DNNs have proven capable of predicting representations across multiple levels of processing. Whereas lower network levels better predict lower level visual representations, subsequent, higher-levels better predict activity in higher-more anterior cortical areas, as measured with functional magnetic resonance imaging (Eickenberg, Gramfort, & Thirion, 2016; Güçlü & van Gerven, 2015; Khaligh-Razavi & Kriegeskorte, 2014; Figure 2B-C). In line with results from macaque IT, DNNs were furthermore able to explain within-category neural similarities, despite being trained on a categorization task that aims at abstracting away from differences across category-exemplars (Khaligh-Razavi & Kriegeskorte, 2014). At a lower spatial, but higher temporal resolution, DNNs have also been shown to be predictive of visually evoked magnetoencephalography (MEG) data (Cichy, Khosla, Pantazis, & Oliva, 2016; Cichy, Khosla, Pantazis, Torralba, & Oliva, 2016; Fritsche, G, Schoffelen, Bosch, & Gerven, 2017). On the behavioural level, deep networks exhibit similar behaviour to humans (Hong, Yamins, Majaj, & DiCarlo, 2016; Kheradpisheh, Ghodrati, Ganjtabesh, & Masquelier, 2016b, 2016a; Kubilius, Bracci, & Op de Beeck, 2016; Majaj, Hong, Solomon, & DiCarlo, 2015) and are currently the best-performing model in explaining human eye-movements in free viewing paradigms (Kümmerer, Theis, & Bethge, 2014). Despite these advances, however, current DNNs still exhibit substantial differences in how they process and recognize visual stimuli (Linsley, Eberhardt, Sharma, Gupta, & Serre, 2017; Rajalingham et al., 2018; Ullman, Assif, Fetaya, & Harari, 2016), how they generalize to atypical category instances (Saleh, Elgammal, & Feldman, 2016), and how they perform under image manipulations, including reduced contrast and additive noise (Geirhos et al., 2017). Yet, the overall success clearly illustrates the power of DNN models for computational neuroscience.

## How can deep neural networks be tested with brain and behavioural data?

DNNs are often trained to optimize external task objectives rather than being derived from neural data. However, even human-level performance does not imply that the underlying computations employ the same mechanisms (Ritter, Barrett, Santoro, & Botvinick, 2017). Testing models with neural measurements is therefore crucial to assess how well network-internal representations match cortical responses. Fortunately, computational neuroscience has a rich toolbox at its disposal that allows researchers to probe even highly complex models, including DNNs (Diedrichsen & Kriegeskorte, 2017).

One such tool are encoding models, which use external, fixed feature spaces in order to model neural responses across a large variety of experimental conditions (e.g. different stimuli, Figure 2A-B). The underlying idea is that if the model and the brain compute the same features, then linear combinations of the model features should enable successful prediction of the neural responses for independent experimental data (Naselaris, Kay, Nishimoto, & Gallant, 2011). For visual representations, the model feature space can be derived from simple filters, such as Gabor-wavelets (Kay, Naselaris, Prenger, & Gallant, 2008), from human labelling of the stimuli (Huth, Nishimoto, Vu, & Gallant, 2012; Mitchell et al., 2008; Naselaris, Prenger, Kay, Oliver, & Gallant, 2009), or from responses in different layers of a DNN (Agrawal, Stansbury, Malik, & Gallant, 2014; Güçlü & van Gerven, 2015).

Probing the system on the level of multivariate response patterns, representational similarity analysis (RSA, Kriegeskorte & Kievit, 2013; Kriegeskorte, Mur, & Bandettini, 2008; Nili et al., 2014) provides another approach to comparing internal representations in DNNs and the brain (Figure 2C). RSA is based around the concept of a representational dissimilarity matrix (RDM), which stores the dissimilarities of a system’s responses (neural or model) to all pairs of experimental conditions. RDMs can therefore be interpreted as describing representational geometries: conditions that elicit similar responses are close together in response space, whereas conditions that lead to differential responses will have larger distances. A model representation is considered similar to a brain representation to the degree that it emphasizes the same distinctions among the stimuli, i.e. model and brain are considered similar, if they elicit similar RDMs. Comparisons on the level of RDMs side-step the problem of defining a correspondence mapping between the units of the model and the channels of brain-activity measurement. This approach can be applied from voxels in fMRI, (Carlin, Calder, Kriegeskorte, Nili, & Rowe, 2011; Guntupalli, Wheeler, & Gobbini, 2016; Khaligh-Razavi & Kriegeskorte, 2014; Kietzmann, Swisher, König, & Tong, 2012), to single-cell recordings (Kriegeskorte et al., 2008; Leibo, Liao, Freiwald, Anselmi, & Poggio, 2017; Tsao, Moeller, & Freiwald, 2008), M/EEG data (Cichy, Pantazis, & Oliva, 2014; Kietzmann, Gert, Tong, & König, 2017), and behavioural measurements including perceptual judgments (Mur et al., 2013).

Although the internal features in a model and the brain may be similar, the distribution of features may not parallel the neural selectivity observed in neuroimaging data. This can either be due to methodological limitations of the neuroimaging technique, or because respective brain area exhibits a bias for certain features that is not captured in the model. To account for such deviations, mixed RSA provides a technique to recombine model features to best explain the empirical data (Khaligh-Razavi, Henriksson, Kay, & Kriegeskorte, 2017). The increase in explanatory power due to this reweighting thereby directly speaks to the question in how far the original, non-reweighted feature space contained the correct feature distribution, relative to the brain measurements.

On the behavioural level, recognition performance (Cadieu et al., 2014; Hong et al., 2016; Majaj et al., 2015; Rajalingham et al., 2018), perceptual confusions, and illusions provide valuable clues as to how representations in brains and DNNs may differ. For instance, it can be highly informative to understand the detailed patterns of errors (Walther, Caddigan, Fei-Fei, & Beck, 2009) and reaction times across stimuli, which may reveal subtle functional differences between systems that exhibit the same overall level of task performance. Visual metamers (Freeman & Simoncelli, 2011; T. S. A. Wallis, Bethge, & Wichmann, 2016) provide a powerful tool to test for similarities in internal representations across systems. Given an original image, a modified version is created that nevertheless leads to an unaltered model response (for instance, the activation profile of a DNN layer). For instance, if a model was insensitive to a selected band of spatial frequencies, then modifications in this particular range will remain unnoticed by the model. If the human brain processed the stimuli via the same mechanism as the model, it should similarly be insensitive to such changes. The two images are therefore indistinguishable (“metameric”) to the model and the brain. Conversely, an adversarial example is a minimal modification of an image that elicits a different category label from a DNN (I. J. Goodfellow, Shlens, & Szegedy, 2015; Nguyen, Yosinski, & Clune, 2015). For convolutional feedforward networks, minimal changes to an image (say of a bus), which are imperceptible to humans, lead the model to classify the image incorrectly (say as an ostrich). Adversarial examples can be generated using the backpropagation algorithm down to the level of the image, to find the gradients in image space that change the classification output. This method requires omniscient access to the system, making it impossible to perform a fair comparison with biological brains, which might likewise be confused by stimuli designed to exploit the idiosyncratic aspects (Elsayed et al., 2018; Kriegeskorte, 2015). The more general lesson for computational neuroscience is that metamers and adversarial examples provide methods for designing stimuli for which different representations disagree maximally. This can optimise the power to adjudicate between alternative models experimentally.

Ranging across levels of description and modalities of brain-activity measurement, from responses in single neurons, to array recordings, fMRI and MEG data, and behaviour, the methods described here enable computational neuroscientists to investigate the similarities and differences between models and neural responses. This essential element is required to be able to find an answer to the question which biological detail and set of computational objectives is needed to align the internal representations of brains and DNNs, while exhibiting successful task-performance.

## Drawing insights from deep neural network models

Deep learning has transformed machine learning and only recently found its way back into computational neuroscience. Despite their high performance in terms of predicting held-out neural data, DNNs have been met with scepticism regarding their explanatory value as models of brain information processing (e.g. Kay, 2017). One of the arguments commonly put forward is that DNNs merely exchange one impenetrably complex system with another (the “black box” argument). That is, while DNNs may be able to predict neural data, researchers now face the problem of understanding what exactly the network is doing.

The black box argument is best appreciated in historical context. Shallow models are easier to understand and supported by stronger mathematical results. For example, the weight template of a linear-nonlinear model can be directly visualised and understood in relation to the concept of an optimal linear filter. Simple models can furthermore enable researchers to understand the role of each individual parameter. A model with fewer parameters is therefore considered more parsimonious as a theoretical account. It is certainly true that simpler models should be preferred over models with excessive degrees of freedom. Many seminal explanations in neuroscience have been derived from simple models. This argument only applies, however, if the two models provide similar predictive power. Models should be as simple as possible, but no simpler. Because the brain is a complex system with billions of parameters (presumably containing the domain knowledge required for adaptive behaviour) and complex dynamics (which implement perceptual inference, cognition, and motor control), computational neuroscience will eventually need complex models. The challenge for the field is therefore to find ways to draw insight from them. One way is to consider their constraints at a higher level of abstraction. The computational properties of DNNs can be understood as the result of four manipulable elements: the *network architecture*, the *input statistics*, the *functional objective*, and the *learning algorithm*.

### Insights generated at a higher-level of abstraction: experiments with network architecture, input statistics, functional objective, and the learning algorithm

A worthwhile thought experiment for neuroscientists is to consider what cortical representations would develop if the world were different. Governed by different input statistics, a different distribution of category occurrences or different temporal dependency structure, the brain and its internal representations may develop quite differently. Knowledge of how it would differ can provide us with principal insights into the objectives that it tries to solve. Deep learning allows computational neuroscientists to make this thought experiment a simulated reality. Investigations of which aspects of the simulated world are crucial to render the learned representations more similar to the brain thereby serve an essential function.

In addition to changes in input statistics, the network architecture can be subject to experimentation. Current DNNs derive their power from bold simplifications. Although complex in terms of their parameter count, they are simple in terms of their component mechanisms. Starting from this abstract level, biological details can be integrated in order to see which ones prove to be required, and which ones do not, for predicting a given neural phenomenon. For instance, it can be asked whether neural responses in a given paradigm are best explained by a feed-forward or a recurrent network architecture. Biological brains draw from a rich set of dynamical primitives. It will therefore be interesting to see to what extent incorporating more biologically inspired mechanisms can enhance the power of DNNs and their ability to explain neural activity and animal behaviour.

Given input statistics and architecture, the missing determinants that transform the randomly initialised model into a trained DNN are the objective function and the learning algorithm. The idea of normative approaches is that neural representations in the brain can be understood as being optimized with regard to one or many overall objectives. These define what the brain should compute, in order to provide the basis for successful behaviour. While experimentally difficult to investigate, deep learning trained on different objectives allows researchers to ask the directly related inverse question: what functions need to be optimized such that the resulting internal representations best predict neural data? Various objectives have been suggested in both the neuroscience and machine learning community. Feed-forward convolutional DNNs are often trained with the objective to minimize classification error (Krizhevsky et al., 2012; Simonyan & Zisserman, 2015; Yamins & DiCarlo, 2016). This focus on classification performance has proven quite successful, leading researchers to observe an intriguing correlation: classification performance is positively related to the ability to predict neural data (Khaligh-Razavi & Kriegeskorte, 2014; Yamins et al., 2014). That is, the better the network performed on a given image set, the better it could predict neural data, even though the latter was never part of the training objective. Despite its success, the objective to minimize classification error in a DNN for visual object recognition requires millions of labelled training images. Although the finished product, the trained DNN, provides the best current predictive model of ventral stream responses, the process by which the model is obtained is not biologically plausible.

To address this issue, additional objective functions from the unsupervised domain have been suggested, allowing the brain (and DNNs) to create error signals without external feedback. One influential suggestion is that neurons in the brain aim at an efficient sparse code, while faithfully representing the external information (Olshausen & Field, 1996; Simoncelli & Olshausen, 2001). Similarly, compression-based objectives aim to represent the input with as few neural dimensions as possible. Autoencoders are one model class following this coding principle (Hinton & Salakhutdinov, 2006). Exploiting information from the temporal domain, the temporal stability or slowness objective is based on the insight that latent variables that vary slowly over time are useful for adaptive behaviour. Neurons should therefore detect the underlying, slowly changing signals, while disregarding fast changes likely due to noise. This potentially simplifies readout from downstream neurons (Berkes & Wiskott, 2005; Földiák, 1991; C. Kayser, Körding, & König, 2003; Christoph Kayser, Einhäuser, Dümmer, König, & Körding, 2001; Körding, Kayser, Einhäuser, & König, 2004; Rolls, 2012; Wiskott & Sejnowski, 2002). Stability can be optimized across layers in hierarchical systems, if each subsequent layer tries to find an optimally stable solution from the activation profiles in previous layer. This approach was shown to lead to invariant codes for object identity (Franzius, Wilbert, & Wiskott, 2008) and viewpoint-invariant place-selectivity (Franzius, Sprekeler, & Wiskott, 2007; Wyss, König, & Verschure, 2006). Experimental evidence in favour of the temporal stability objective in the brain has been provided by electrophysiological and behavioural studies (N. Li & DiCarlo, 2008, 2010; G. Wallis & Bülthoff, 2001).

Many implementations of classification, sparseness and stability objectives ignore the action repertoire of the agent. Yet, different cognitive systems living in the same world may exhibit different neural representations because the requirements to optimally support action may differ. Deep networks optimizing the predictability of the sensory consequence (Weiller, Märtin, Dähne, Engel, & König, 2010), or cost of a given action (Mnih et al., 2015) have started incorporating the corresponding information. On a more general note, it should be noted that there are likely multiple objectives that the brain optimizes across space and time (Marblestone, Wayne, & Kording, 2016), and neural response patterns may encode multiple types of information simultaneously, enabling selective read-out by downstream units (DiCarlo & Cox, 2007).

In summary, one way to draw theoretical insights from DNN models is to explore what architectures, input statistics, objective functions, and learning algorithms yield the best predictions for neural activity and behaviour. This approach does not elucidate the role of individual units or connections in the brain. However, it can reveal what features of biological structure likely support selected functional aspects, and what objectives the biological system might be optimised for, either via evolutionary pressure, or during the development of the individual.

### Looking into the black box: receptive field visualization and “in silico” electrophysiology

In addition to contextualising DNNs on a more abstract level, we can also open the ‘black box’ and look inside. Unlike a biological brain, a DNN model is entirely accessible to scrutiny and manipulation, enabling, for example, high-throughput “in silico” electrophysiology. The latter can be used to gain an intuition for the selectivity of individual units. For instance, large and diverse image sets can be searched for the stimuli that lead to maximal unit activation (Figure 3). Building on this approach, the technique of network dissection has emerged, which provides a more quantitative view on unit selectivity (Zhou, Bau, Oliva, & Torralba, 2017). It uses a large dataset of segmented and labelled stimuli to first find images and image regions that maximally drive network units. Based on the ground-truth labels for these images, it is then derived whether the unit’s selectivity is semantically consistent across samples. If so, an interpretable label, ranging from colour-selectivity, to different textures, object parts, objects, and whole scenes, is assigned to the unit. This characterization can be applied to all units of a network layer, providing powerful summary statistics.

**Figure 3.**
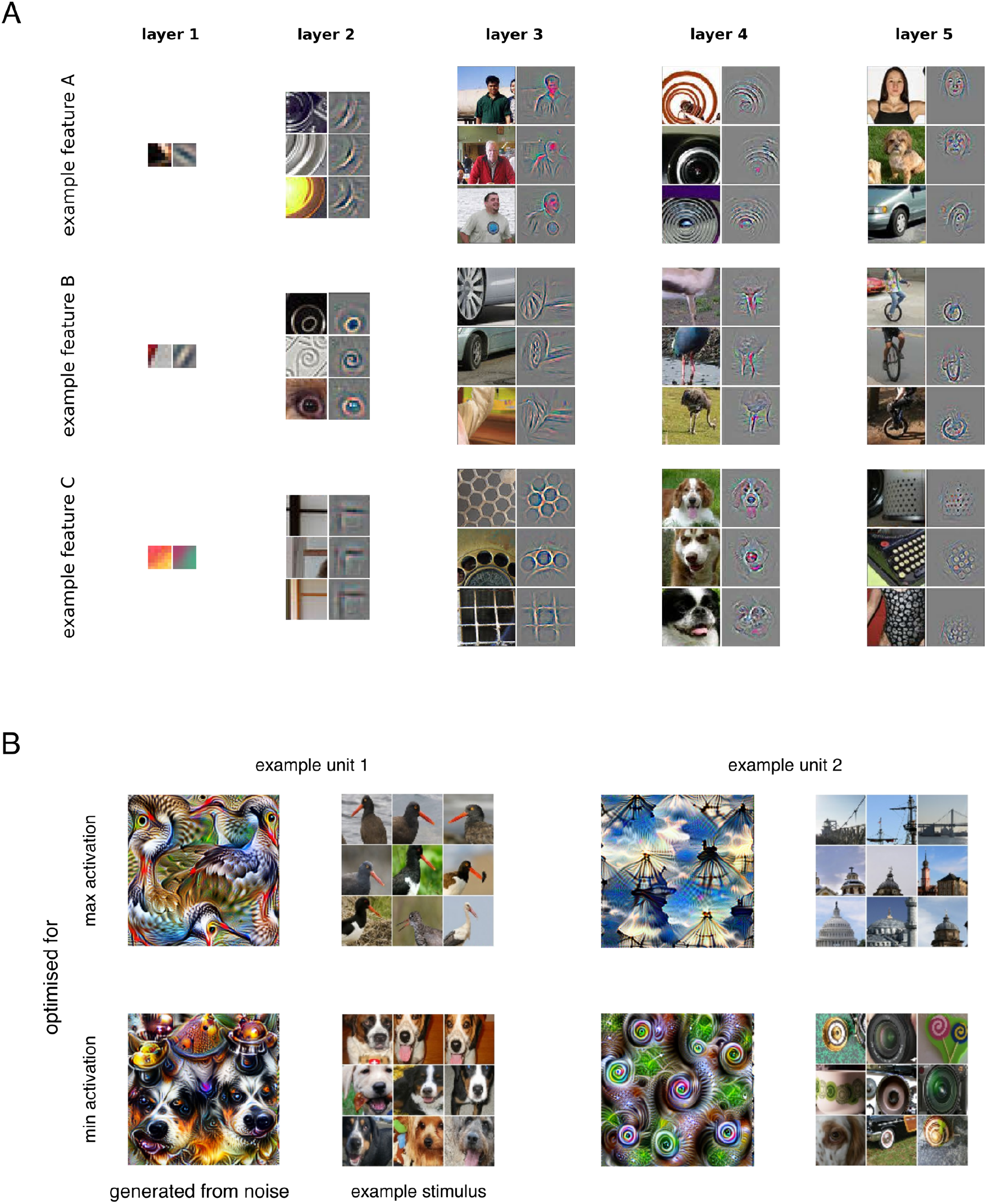
Visualizing the preferred features of internal neurons. (A) Activations in a random subset of feature maps across layers for strongly driving ImageNet images projected down to pixel space (adapted from Zeiler & Fergus, 2014). (B) Feature visualization based on image optimization for two example units. Starting from pure noise, images were altered to maximally excite, or inhibit the respective network unit. Maximally and minimally driving example stimuli are shown next to the optimization results (adapted from Olah et al., 2017).

Another method for understanding a unit’s preferences is via feature visualization, a rapidly expanding set of diverse techniques that directly speak to the desire for human interpretability beyond example images. One of many ways to visualize what image features drive a given unit deep in a neural network is to approximately undo the operations performed by a convolutional DNN in the context of a given image (Zeiler & Fergus, 2014). This results in visualisations such as those shown in Figure 3A. A related technique is feature visualisation by optimization (see Olah, Mordvintsev, & Schubert (2017) for a review), which is based on the idea to use backpropagation (Rumelhart, Hinton, & Williams, 1986) potentially including a natural image prior, to calculate the change in the input needed to drive or inhibit the activation of any unit in a DNN (Simonyan & Zisserman, 2015; Yosinski, Clune, Nguyen, Fuchs, & Lipson, 2015). As one option, the optimisation can be started from an image that already strongly drives the unit, computing a gradient in image space that enhances the unit’s activity even further. The gradient-adjusted image shows how small changes to the pixels affect the activity of the unit. For example, if the image that is strongly driving the unit shows a person next to a car, the corresponding gradient image might reveal that it is really the face of the person driving the unit’s response. In that case, the gradient image would deviate from zero only in the region of the face and adding it to the original image would accentuate the facial features. Relatedly, optimisation can be started from an arbitrary image, with the goal of enhancing the activity of a single or all units in a given layer (as iteratively performed in Google’s DeepDream). Another option is to start from pure noise images, and to again use backpropagation to iteratively optimise the input to strongly drive a particular unit. This approach yields complex psychedelic looking patterns containing features and forms, which the network has learned through its task training (Figure 3B). Similar to the previous approach that characterizes a unit by finding maximally driving stimuli, gradient images are best derived from many different test images to get a sense of the orientation of its tuning surface around multiple reference points (test images). Relatedly, it is important to note that the tuning function of a unit deep in a network cannot be characterised by a single visual template. If it could, there would be no need for multiple stages of nonlinear transformation. However, the techniques described in this section can provide first intuitions about unit selectivities across different layers or time-points.

DNNs can provide computational neuroscientists with a powerful tool, and are far from a black box. Insights can be generated by looking at the parameters of DNN models at a more abstract level. For instance, by observing the effects on predictive performance resulting from changes to the network architecture, input statistics, objective function, and learning algorithm. Furthermore, in silico electrophysiology enables researchers to measure and manipulate every single neuron, in order to visualize and characterize its selectivity and role in the overall system.

## What neurobiological details matter to brain computation?

A second concern about DNNs is that they abstract away too much from biological reality to be of use as models for neuroscience. Whereas the black box argument states that DNNs are too complex, the biological realism argument states that they are too simple. Both arguments have merit. It is conceivable that a model is simultaneously too simple (in some ways) and too complex (in other ways). However, this raises a fundamental question: Which features of the biological structure should be modelled and which omitted to explain brain function (Tank, 1989)?

Abstraction is the essence modelling and is the driving force of understanding. If the goal of computational neuroscience is to understand brain *computation*, then we should seek the simplest models that can explain task performance and predict neural data. The elements of the model should map onto the brain at some level of description. However, what biological elements must be modelled is an empirical question. DNNs are important not because they capture many biological features, but because they provide a minimal functioning starting point for exploring what biological details matter to brain computation. If, for instance, spiking models outperformed rate-coding models at explaining neural activity and task performance (for example in tasks requiring probabilistic inference (Buesing, Bill, Nessler, & Maass, 2011)), then this would be strong evidence in favour of spiking models. Large-scale models will furthermore enable an exploration of the level of detail required in systems implementing the whole perception-action cycle (Eliasmith et al., 2012; Eliasmith & Trujillo, 2014).

Convolutional DNNs like AlexNet (Krizhevsky et al., 2012), and VGG (Simonyan & Zisserman, 2015) were built to optimise performance, rather than biological plausibility. However, these models draw from a history of neuroscientific insight and share many qualitative features with the primate ventral stream. The defining property of convolutional DNNs is the use of convolutional layers. These have two main characteristics: (1) local connections that define receptive fields and (2) parameter sharing between neurons across the visual field. Whereas spatially restricted receptive fields are a prevalent biological phenomenon, parameter sharing is biologically implausible. However, biological visual systems learn qualitatively similar sets of basis features in different parts of a retinotopic map, and similar results have been observed in models optimizing a sparseness objective (Güçlü & van Gerven, 2014; Olshausen & Field, 1996). Moving toward greater biological plausibility with DNNs, locally connected layers that have receptive fields without parameter sharing were suggested (Uetz & Behnke, 2009). Researchers have already started exploring this type of DNN, which was shown to be very successful in face recognition (Sun, Wang, & Tang, 2015; Taigman, Ranzato, Aviv, & Park, 2014). One reason for this is that locally connected layers work best in cases where similar features are frequently present in the same visual arrangement, such as faces. In the brain, retinotopic organization principles have been proposed for higher-level visual areas (Levy, Hasson, Avidan, Hendler, & Malach, 2001), and similar organisation mechanisms may have led to faciotopy, the spatially stereotypical activation for facial features across the cortical surface in face-selective regions (Henriksson, Mur, & Kriegeskorte, 2015).

### Beyond the feed-forward sweep: recurrent DNNs

Another aspect in which convolutional AlexNet and VGG deviate from biology is the focus on feed-forward processing. Feedforward DNNs compute static functions, and are therefore limited to modelling the feed-forward sweep of signal flow through a biological system. Yet, recurrent connections are a key computational feature in the brain, and represent a major research frontier in neuroscience. In the visual system, too, recurrence is a ubiquitous phenomenon. Recurrence is likely the source of representational transitions from global to local information (Matsumoto, Okada, Sugase-Miyamoto, Yamane, & Kawano, 2005; Sugase, Yamane, Ueno, & Kawano, 1999). The timing of signatures of facial identity (Barragan-Jason, Besson, Ceccaldi, & Barbeau, 2013; Freiwald & Tsao, 2010) and social cues, such as direct eye-contact (Kietzmann et al., 2017), too, point towards a reliance on recurrent computations. Finally, recurrent connections likely play a vital role in early category learning (Kietzmann, Ehinger, Porada, Engel, & König, 2016), in dealing with occlusion (Oord, Kalchbrenner, & Kavukcuoglu, 2016; Spoerer, McClure, & Kriegeskorte, 2017; Wyatte, Curran, & O’Reilly, 2012; Wyatte, Jilk, & O’Reilly, 2014) and object-based attention (Roelfsema, Lamme, & Spekreijse, 1998).

Whereas the first generation of DNNs focused on feed-forward, the general class of DNNs can implement recurrence. By using lateral recurrent connections, DNNs can implement visual attention mechanisms (Z. Li, Yang, Liu, Wen, & Xu, 2017; Mnih, Heess, Graves, & Kavukcuoglu, 2014), and lateral recurrent connections can also be added to convolutional DNNs (Liang & Hu, 2015; Spoerer et al., 2017). These increase the effective receptive field size of each unit, and allow for long-range activity propagation (Pavel et al., 2017). Lateral connections can make decisive contributions to network computation. For instance, in modelling the responses of retinal ganglion cells, the introduction of lateral recurrent connections to feed-forward CNNs lead to the emergence of contrast adaptation in the model (McIntosh, Maheswaranathan, Nayebi, Ganguli, & Baccus, 2017). In addition to local feedforward and lateral recurrent connections, the brain also uses local feedback, as well as long-range feedforward and feedback connections. While missing from the convolutional DNNs previously used to predict neural data, DNNs with these different connection types have been implemented (He et al., 2016; Liao & Poggio, 2016; Srivastava, Greff, & Schmidhuber, 2015). Moreover, long short-term memory (LSTM) units (Hochreiter & Schmidhuber, 1997) are a popular form of recurrent connectivity used in DNNs. These units use differentiable read and write gates to learn how to use and store information in an artificial memory “cell”. Recently, a biologically plausible implementation of LSTM units has been proposed using cortical microcircuits (Costa, Assael, Shillingford, de Freitas, & Vogels, 2017).

The field of recurrent convolutional DNNs is still in its infancy, and the effects of lateral and top-down connections on the representational dynamics in these networks, as well as their predictive power for neural data are yet to be fully explored. Recurrent architectures are an exciting tool for computational neuroscience and will likely allow for key insights into the recurrent computational dynamics of the brain, from sensory processing to flexible cognitive tasks (Song, Yang, & Wang, 2016, 2017).

### Optimising for external objectives: backpropagation and biological plausibility

Apart from architectural considerations, backpropagation, the most successful learning algorithm for DNNs, has classically been considered neurobiologically implausible. Rather than as a model of biological learning, backpropagation may be viewed as an efficient way to arrive at reasonable parameter estimates, which are then subject to further tests. That is, even if backpropagation is considered a mere technical solution, the trained model may still be a good model of neural system. However, there is also a growing literature on biologically plausible forms of error-driven learning. If the brain does optimise cost functions during development and learning (which can be diverse, and supervised, unsupervised, or reinforcement-based), then it will have to use a form of optimization mechanism, an instance of which are stochastic gradient descent techniques. The current literature suggests several neurobiologically plausible ways in which the brain could adjust its internal parameters to optimise such objective functions (Lee, Zhang, Fischer, & Bengio, 2015; Lillicrap et al., 2016; O’Reilly, 1996; Xie & Seung, 2003). These methods have furthermore been shown to allow deep neural networks to learn simple vision tasks (Guerguiev, Lillicrap, & Richards, 2017). The brain might not be performing the exact algorithm of backpropagation, but it might have a mechanism for modifying synaptic weights in order to optimise one or many objective functions (Marblestone et al., 2016).

### Stochasticity, oscillations, and spikes

Another aspect in which DNNs deviate from biological realism is that DNNs are generally deterministic, while biological networks are stochastic. While much of this stochasticity is commonly thought to be noise, it has been hypothesized that this variability could code for uncertainty (Fiser, Berkes, Orbán, & Lengyel, 2010; Hoyer, Hyvarinen, Patrik, Aapo, & Hyv, 2003; Orban, Berkes, Fiser, & Lengyel, 2016). In line with this, DNNs that include stochastic sampling during training and test can yield higher performance, and are better able to estimate their own uncertainty (McClure & Kriegeskorte, 2016). Furthermore, current recurrent convolutional DNNs often only run for a few time steps, and the roles of dynamical features found in biological networks, such as oscillations, are only beginning to be tested (Finger & König, 2013; Reichert & Serre, 2013; Siegel, Donner, & Engel, 2012). Another abstraction is the omission of spiking dynamics. However, DNNs with spiking neurons can be implemented (Hunsberger & Eliasmith, 2016; Tavanaei & Maida, 2016) and represent an exciting frontier of deep learning research. These considerations show that it would be hasty to judge the merits of DNNs based on the level of abstraction chosen in the first generation.

## Deep learning: a powerful framework to advance computational neuroscience

Deep neural networks have revolutionised machine learning and AI, and have recently found their way back into computational neuroscience. DNNs reach human-level performance in certain tasks, and early experiments indicate that they are capable of capturing characteristics of cortical function that cannot be captured with shallow linear-nonlinear models. With this, DNNs offer an intriguing new framework that enables computational neuroscientists to address fundamental questions about brain computation in the developing and adult brain.

Computational neuroscience comprises a wide range of models, defined at various levels of *biological* and *behavioural detail* (Figure 4). For instance, many conductance-based models contain large amounts of parameters to explain single or few neurons at great level of detail but are typically not geared towards behaviour. DNNs, at the other end of the spectrum, use their high number of parameters not to account for effects on the molecular level, but to achieve behavioural relevance, while accounting for overall neural selectivity. Explanatory merit is not only gained by biological realism (because this would render human brains the perfect explanation for themselves), nor does it directly follow from simplistic models that cannot account for complex animal behaviour. The space of models is continuous and neuroscientific insight works across multiple levels of explanation, following top-down and bottom-up approaches (Craver, 2009). The usage of DNNs in computational neuroscience is still in its infancy, and the integration of biological detail will require close collaboration between modellers, experimental neuroscientists, and anatomists.

**Figure 4.**
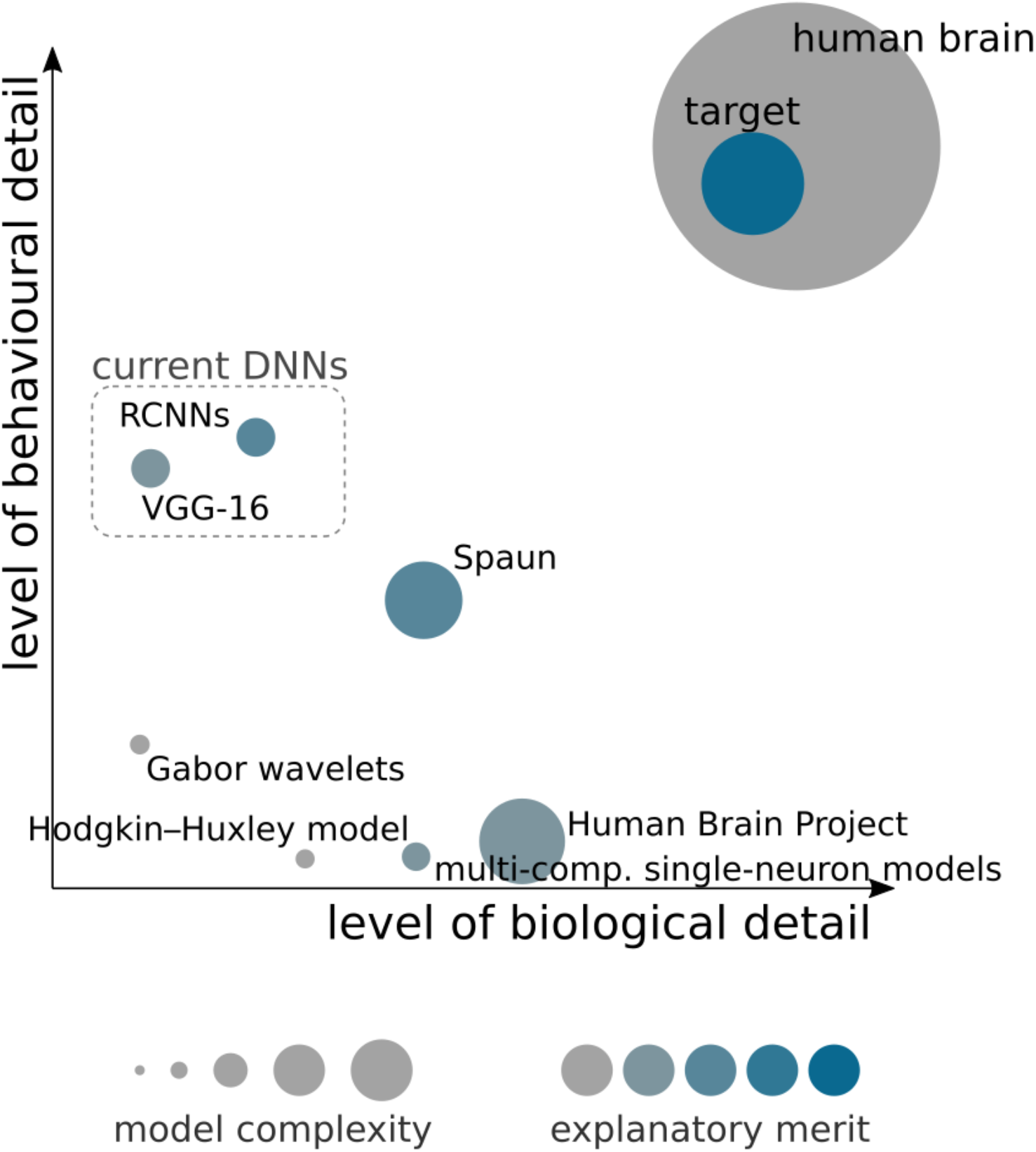
Cartoon overview of different models in computational neuroscience. Given computational constraints, models need to make simplifying assumptions. These can either be regarding the biological detail, or behavioral relevance of the model output. The explanatory merit of a model is not dependent on the exact replication of biological detail, but on its ability to provide insights into the inner workings of the brain at a given level of abstraction.

DNNs will not replace shallow models, but rather enhance the researchers’ investigative repertoire. With computers approaching the brain in computational power, we are entering a truly exciting phase of computational neuroscience.

## Further reading

- Kriegeskorte (2015) – introduction of deep learning as a general framework to understand brain information processing
- Yamins, & DiCarlo (2016) – perspective on goal-driven deep learning to understand sensory processing
- Marblestone et al. (2016) – review with a focus on cost functions in the brain and DNNs
- Lindsay (2018) – overview of how DNNs can be used as models of visual processing
- LeCun et al. (2015) – high level overview of deep learning developments
- Googfellow et al. (2016) – introductory book on deep learning

